# Predictor species: Improving assessments of rare species occurrence by modelling environmental co-responses

**DOI:** 10.1101/546879

**Authors:** Peter R. Thompson, William F. Fagan, Phillip P.A. Staniczenko

## Abstract

Designing an effective conservation strategy requires understanding where rare species are located. Although species distribution models are primarily used to identify patterns at large spatial scales, their general methodology is relevant for predicting the occurrence of individual species at specific locations. Here we present a new approach that uses Bayesian networks to improve predictions by modelling environmental co-responses among species. For species from a European peat bog community, our approach consistently performs better than single-species models, and better than conventional multi-species models for rare species when calibration data are limited. Furthermore, we identify a group of “predictor species” that are relatively common, insensitive to the presence of other species, and can be used to improve occurrence predictions of rare species. Predictor species are distinct from other categories of conservation surrogates such as umbrella or indicator species, which motivates focused data collection of predictor species to enhance conservation practices.

## INTRODUCTION

Species distribution models (SDMs) are widely used in ecology to predict the geographical ranges of individual species^1–5^, and multiple SDMs can be interpreted together to estimate the composition of an ecological community at a particular location^6–8^. SDMs are also used to aid in the conservation of rare species that occur at relatively few locations compared to other species in the community^9–11^. Because rare species often have specialized habitat preferences^12^ and are harder to detect^13^, protecting areas where rare species are known to occur or, more realistically, are expected to occur, is critical for preserving the Earth’s biodiversity^14^. However, protecting the wrong areas due to model inaccuracy is a costly mistake that does little to promote the survival of rare and threatened species^15^.

The growing desire and potential for SDMs to make predictions at smaller spatial scales has led to an integration of ideas from macroecology and community ecology^16,17^. Ecologists initially made predictions using environment-only SDMs that included only abiotic variables like temperature and rainfall^18^, but soon recognized that incorporating dependencies among species was necessary to explain empirical distribution patterns^19–24^. Recent work has explored a variety of approaches to modelling such dependencies in SDMs^25–29^, and a simple yet successful strategy involves including the presence or absence of non-focal species as independent variables in generalized linear models (GLMs)^30,31^ and maximum entropy models^32^. However, this strategy has not always improved results; for example, predictions for rare species from a British plant community were less accurate with multi-species models than with single-species versions of two machine-learning methods^33^. A more comprehensive approach to modelling shared environmental co-responses involves joint species distribution models^34,35^, but calibrating these models requires species co-occurrence data that can be time-consuming and labor-intensive to collect. Bayesian networks (BNs) offer a balanced approach to modelling how the presence of a focal species is affected by the presence or absence of other species^17^. When applied to an ecological community, a BN adjusts “prior” probabilities of species occurrence from environment-only models to produce “posterior” probabilities that also reflect the effect of biotic interactions and other interspecific relationships among species.

Here our goal is to improve assessments of species occurrence at specific locations, especially for rare species, by including information on environmental responses in SDM-like predictive models. We compare the performance of three types of model: (i) environment-only GLMs (“eGLM”); (ii) multi-species GLMs that include the presence or absence of non-focal species as additional independent variables (“sGLM”); and (iii) a new approach that combines probabilities from the eGLM with a BN that represents strong environmental co-responses among species (“eGLM+BN”). We compare these three models to an approach based on joint species distribution modelling that provides an upper bound to model accuracy because it requires much more input data for calibration. We test models using data on 54 plant species from a European peat bog community at 56 locations^36^. We find that the two multi-species models consistently outperform the eGLM, with the eGLM+BN able to produce accurate predictions even with limited calibration data. Based on a BN for the peat bog community, we identify a group of “predictor species” that are useful for improving predictions of rare species occurrence. We suggest that predictor species could function as conservation surrogates if a stated aim is to establish geographical distributions of rare species. To this end, predictor species complement existing categories of conservation surrogates such as umbrella species (typically found at many locations^37^) and indicator species (typically found at locations with high species richness^38^), which are less suitable for informing spatial distributions of regional biodiversity^39^.

## METHODS

### Data

We tested our approach using data on a peat bog community of 54 plant species at 56 locations across Europe^36^. Data included abundance records for each plant species and bioclimate data for each location. Of nine available bioclimate variables, we included four in generalized linear models: mean annual temperature, mean annual precipitation, latitude and temperature seasonality (seasonality being the difference in a property between the warmest and coolest month in a given year). These four variables had the highest average correlations with species occurrence (Appendix S1: Table S1) and were not highly correlated with each other (Appendix S1: Table S2). Because our goal was to develop models for predicting the occurrence of individual species at specific locations, we converted species abundances at each location to a binary presence-absence measure (any species with an abundance over 0 was considered to be present) to use as a dependent variable for calibrating and testing models.

### Modelling occurrence predictions using only environmental variables (eGLM)

We used generalized linear models^40,41^ to create environment-only predictions for the plant species in the peat bog community. The eGLM included only bioclimate data as independent variables, with the presence or absence of the focal species at a specific location as the dependent variable:

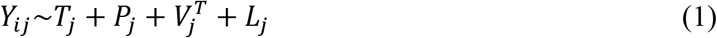

where *Y_ij_* is the presence or absence of species *i* at location *j*; and *T_j_* is mean annual temperature, *P_j_* is mean annual precipitation, 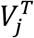 is temperature seasonality, and *L_j_* is latitude, at location *j*. We used a logit link function between independent and dependent variables.

### Estimating environmental co-responses among species

To develop models that incorporated the occurrence of non-focal species into co-occurrence predictions, we constructed a correlation matrix outlining the strength of all the interspecific relationships in the peat bog community. First, we computed the Pearson correlation between the presence or absence of each pair of species across the 56 locations. The result was a symmetric 54-by-54 species correlation matrix with ones on the leading diagonal. We then set the ones on the leading diagonal to 0 and specified a threshold value to convert all off-diagonal entries to 0, 1, or −1, depending on whether the correlation was above the threshold value and positive or negative. We used 0.35 as the threshold value because it represented a point of inflection in the resulting number of nonzero entries in the transformed correlation matrix (Appendix S1: Fig. S1). The transformed correlation matrix had a total of 184 nonzero entries (130 positive and 54 negative), and only 7 of the 54 species did not have a non-zero entry with any other species in the community.

### Modelling environmental co-responses among species as independent variables (sGLM)

The sGLM included the occurrence of non-focal species as additional independent variables:

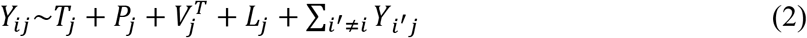

where the final summation term only includes species that have been shown to strongly influence the occurrence of species i according to the correlation matrix (note that each non-focal species *i*’ has a unique GLM slope coefficient), which ensured that the sGLM describes the same environmental co-responses as the eGLM+BN, see below. For species without any such influences, the eGLM, sGLM, and eGLM+BN all give identical results.

### Modelling environmental co-responses among species using a Bayesian network (eGLM+BN)

A BN represents environmental co-responses as conditional dependencies between the occurrence probabilities of individual species in a community^17^. Compared to some multi-species models that include the occurrence of non-focal species as additional independent variables (e.g., sGLM), the BN is applied as a separate, secondary step after environment-only models. We based the BN in the present study on the above correlation matrix of environmental co-responses among species. In this application, occurrence probabilities from the eGLM, so-called “prior” probabilities, are combined with the BN to obtain “posterior” probabilities that reflect environmental co-responses among species.

The BN must be a directed acyclic graph, meaning that directed edges representing conditional dependencies point from one species to another and, there is no way of returning to a species by following a sequence of directed edges originating from that species^17^. To satisfy these criteria, we implemented a hierarchy for the 54 species such that directed edges point from species higher up in the hierarchy to those lower down. We used a hierarchy based on species abundance (aggregated across the 56 locations), with directed edges pointing from more abundant species to less abundant species. Starting with the transformed correlation matrix, we removed any non-zero entries associated with edges that pointed from a less-to more-abundant species. The result was a BN with 65 positive and 27 negative conditional dependencies involving 47 of the 54 species (Appendix S1: Fig. S2). We used the Boolean “OR” rule to determine how prior probabilities from the eGLM are converted to posterior probabilities when a species has multiple conditional dependencies in the BN^17^ (see Appendix S1: Fig. S3 for a description and worked example of the Boolean “OR” rule).

### Evaluating model performance

We evaluated the effect of data availability on model performance by using a fraction of the empirical data in a training partition to calibrate models and the remaining data in a test partition to measure predictive accuracy. We considered three proportions of training and test partition sizes: 25% training and 75% test, 50% training and 50% test, and 75% training and 25% test. We ran 1000 randomizations of data for each proportion. We measured the predictive accuracy of each model using the area under receiver operating characteristic curve (AUC) method, which measures the ability of an SDM to discriminate between known species presences and absences^42^. We also considered true skill statistic (TSS^43^) but found it resulted in such high variability between randomizations (Appendix S1: Fig. S4) that we were not as easily able to distinguish between models as with AUC.

To obtain an upper bound to model performance, we modified the joint species distribution model (JSDM) proposed by Ovaskainen et al.^34^, which requires all co-occurrence relationships among species in a community to be fully quantified. Our JSDM-inspired approach represents the probability of occurrence of a species as a random variable in a jointly distributed set of normal random variables; that is, co-occurrence relationships between species are described by correlations between random variables. Each component of this multivariate distribution—one univariate normal random variable for one species—is centered at the original eGLM estimate for a species, i.e., with no correlations between random variables this approach reduces to a set of independent eGLMs. At the multivariate level, these correlations are organized into a symmetric correlation matrix containing values between −1 and 1. We used the same correlation matrix described above without any simplification. Because the full correlation matrix is used, the distribution of each component depends on the value of the other components. From a statistical standpoint, this means that we can use the method of conditional probabilities to obtain a revised distribution for each species given the known values of the others^44^. In other words, the probability that a species is present at a particular location requires knowing the occurrences of all other species at that location. The amount of information contained in the JSDM-inspired approach means it is expected to produce very good predictions. But the large amounts of data required for parametrization compared to the eGLM, sGLM and eGLM+BN means its output should be considered a practically unattainable upper bound. The data requirements of each model are summarized in Table S3 of Appendix S1.

### Identifying co-responsive species

We identified a group of species whose occurrence predictions were greatly improved by the addition of the BN. We measured the overall benefit the BN added to environment-only models using ΔAUC, which we defined as the difference in AUC scores between the eGLM and the eGLM+BN for an individual species when data were partitioned into 50% training and 50% test. We ran 10 sets of 100 randomizations, considering species with ΔAUC above 0.08 in at least 9 of the 10 sets to be “co-responsive species” (Appendix S1: Table S4). We used boosted regression tree analysis^45,46^ to investigate the shared properties of co-responsive species. Boosted regression tree analysis assigned a “relative importance” to six species properties according to each property’s ability to explain ΔAUC values among co-responsive species (see Supplementary Information). Relative importance values sum to one for all six variables. The six species properties we considered were: the number of incoming BN edges, the proportion of locations where species occurred (“rarity”), the average abundance at locations where each species occurred, the average eGLM AUC score, whether a species was a vascular plant or a *Sphagnum* moss, and topological importance. Topological importance is a summary statistic used in graph theory to evaluate the contribution of each node (in this case, each species) to the overall connectedness of the graph; it has been used to determine keystone species in ecological communities^47^.

### Identifying predictor species

We identified a group of “predictor species” that had a strong effect on the occurrence probabilities of co-responsive species. We defined predictor species as (i) species with multiple outgoing BN edges directly connected to co-responsive species or (ii) species with any outgoing BN edges directly connected to the first set of predictor species.

We compared this set of predictor species to umbrella^37^ and indicator^38^ species from the peat bog community to gauge the extent of overlap between the three groups. Here, we defined umbrella species as species that occurred at 42 (75%) or more of the 56 locations and indicator species as species that, on average, occurred at locations with at least 20 plant species. We chose 20 as a cut-off because only 15 locations (26.8%) satisfied this criterion.

We measured the collective effect of predictor species by computing AUC scores for the eGLM+BN with a partial BN containing only edges among co-responsive and predictor species. As with the original BN, we ran 1000 samples with 25%, 50% and 75% training data, then compared ΔAUC values between partial and full BNs for each co-responsive species.

## RESULTS

### Predicting species occurrence at specific locations

We found that modelling environmental co-responses with both multi-species models consistently improved predictions of species occurrence relative to the eGLM. The eGLM+BN performed better than the sGLM when fewer data were used for model training, but the sGLM performed better when more data were used for model training (Figure 1). With 25% training data, the eGLM+BN yielded an average AUC score of 0.698, compared to the sGLM average of 0.675 and the eGLM average of 0.668. With 50% training data, the eGLM+BN yielded an average AUC score of 0.741, compared to the sGLM average of 0.730 and the eGLM average of 0.711. With 75% training data, the eGLM+BN yielded an average AUC score of 0.772, compared to the sGLM average of 0.775 and the eGLM average of 0.754. As expected, AUC scores for all models increased in line with the amount of data used for model training. The upper-bound AUC estimates provided by our JSDM-inspired approach were 0.817, 0.844, and 0.848 with 25%, 50% and 75% training data, respectively.

**Figure 1.**
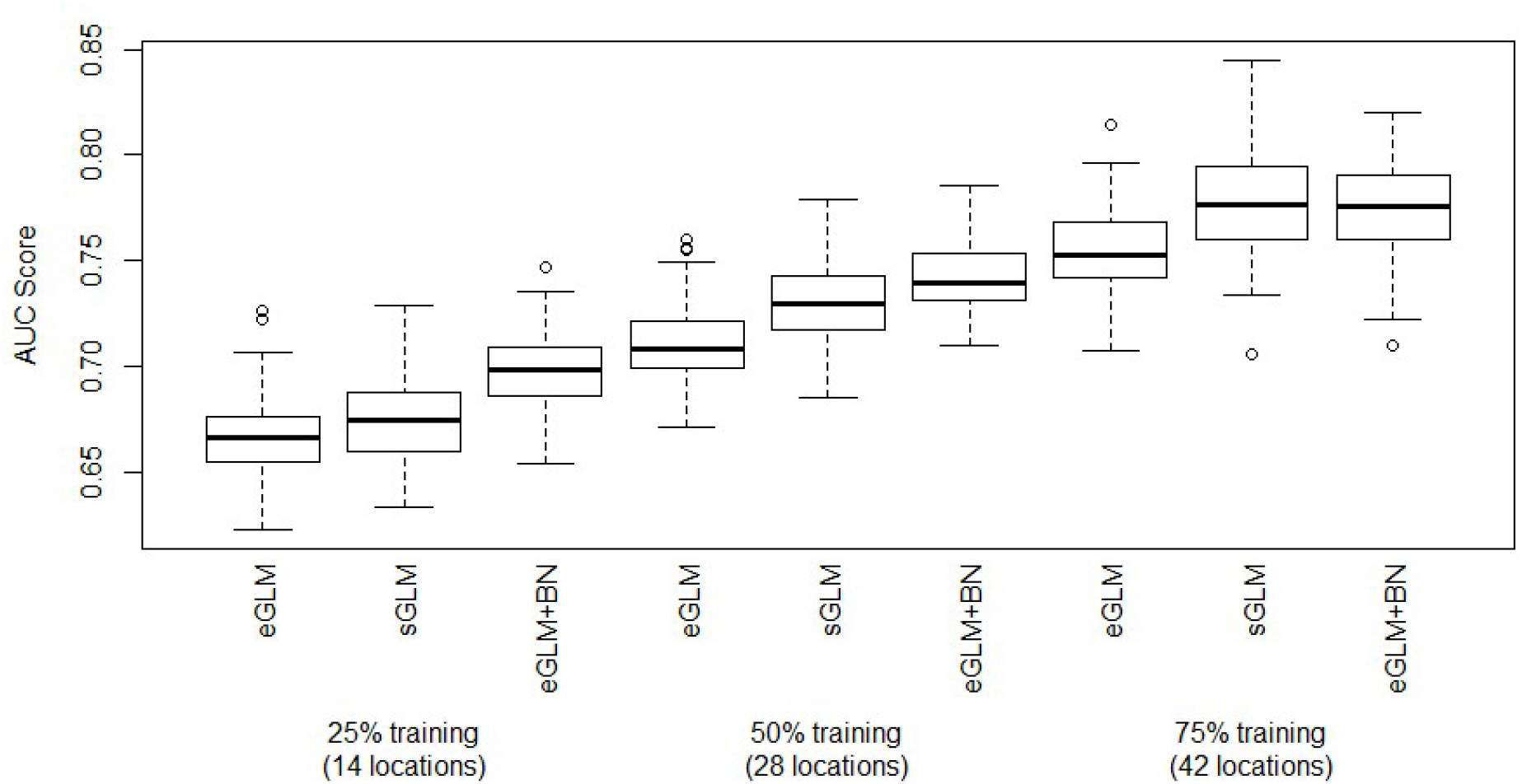
Performance of the eGLM, sGLM, and eGLM+BN measured by AUC at three training partition sizes. The sGLM and eGLM+BN both outperform the eGLM at all partition sizes (1000 random partitions of the 56 locations for each combination of model and training partition size). Notice that the eGLM+BN outperforms the sGLM with 25% training data, but not at 75% training data. With an unrealistic amount of data available for prediction, we observed AUC scores of 0.848 ± 0.042, 0.844 ± 0.032, and 0.817 ± 0.030 at 75%, 50% and 25% training respectively (mean ± standard deviation).

The eGLM+BN improved predictions for almost every species in the peat bog community. We focused further analysis on this model to better understand its increased prediction accuracy with limited amounts of calibration data compared to the eGLM and sGLM. Aside from the 14 species without any incoming BN edges (by definition the BN does not modify predictions for these species), ΔAUC values were positive for all but 6 species; the remaining 40 species had an average ΔAUC value of 0.040 ± 0.041 (mean ± standard deviation), and only 5 of these species had ΔAUC values below 0.01.

### Characterizing co-responsive species whose occurrence patterns are strongly influenced by other species

Of the 54 species from the peat bog community, we identified 6 species with ΔAUC values consistently above 0.08, indicating that the eGLM+BN was particularly effective at improving predictions for these species. We used boosted regression tree analysis^45,46^ to investigate the shared properties of these co-responsive species. We found that rarity had the highest relative importance value of the 6 properties we considered (Table 1). This result suggests that co-responsive species are characterized as being rare—indeed, they occurred at an average of only 11.6% of the 56 locations, compared to the community-wide average of 34.1% (We explored whether this finding may have arisen due to our use of an abundance-based hierarchy to specify the direction of BN edges, but further analysis showed that this choice of hierarchy was not responsible for the result that co-responsive species are typically rare species; see Supplementary Information.). Five of the six co-responsive species were particularly rare (occurring at less than 15% of the 56 peat bog locations). The next most important property was the eGLM AUC average for the species, suggesting that the BN is especially beneficial when environmental variables on their own provide relatively poor predictions of species’ occurrences. The six co-responsive species had an average eGLM AUC of 0.665, compared to the overall average of 0.710.

**TABLE 1.** Relative importance of six properties associated with species according to boosted regression tree analysis.

| Property | Relative Importance (%) |
| --- | --- |
| Rarity | 34.9 |
| Average eGLM AUC score with 50% training data | 24.2 |
| Average abundance | 18.5 |
| Number of incoming BN edges | 15.8 |
| Topological importance | 6.1 |
| *Sphagnum* moss or vascular plant | 0.5 |

### Characterizing predictor species that improve occurrence predictions of other species

We identified 8 predictor species that had a strong effect on the occurrence probabilities of co-responsive species. Two of the predictor species had multiple outgoing BN edges pointing directly to co-responsive species, while the other six indirectly influenced co-responsive species through BN edges with the first predictor species (One of the co-responsive species, *Vaccinium vitis-idea*, actually met the criteria for a predictor species by having two outgoing BN edges pointing towards other co-responsive species, but we chose to consider it only as a co-responsive species in subsequent analysis). Predictor species generally had high eGLM AUC scores and low ΔAUC values. The average eGLM AUC score for predictor species was 0.754 with 50% training data, higher than the overall average of 0.710. Predictor species had an average ΔAUC value of 0.009, lower than the overall average of 0.029 and much lower than the co-responsive species average of 0.114. Taken together, these results suggest that a predictor species is relatively insensitive to the presence or absence of other species and their occurrence is well predicted by abiotic conditions alone. Predictor species were more common than usual but not especially widespread; on average, each predictor species occurred at 45.1% of the 54 locations.

### Analyzing a partial Bayesian network of co-responsive and predictor species

We investigated the performance of a BN containing only edges among co-responsive and predictor species (Figure 2). The partial BN was highly connected with multiple pathways of influence between species. For example, *Sphagnum fallax* (a predictor species) had only one edge pointing directly to a co-responsive species, yet it indirectly influenced five of the six co-responsive species. The partial BN generally yielded better AUC scores than the original BN at all three training partition sizes, despite the partial BN retaining only 19 (12 positive and 7 negative) of the 92 edges in the original BN (including only 9 of 17 edges pointing directly to co-responsive species). Compared to the original BN, which produced ΔAUC values of 0.117 ± 0.065 (mean ± standard deviation) for the co-responsive species, the partial BN produced ΔAUC values of 0.147 ± 0.068 (Appendix S1: Table S5). Compared to the original BN, the reduced nature of the partial BN made species occurrence probabilities much easier to compute, while also lowering variability and noise caused by unnecessary BN edges.

**Figure 2.**
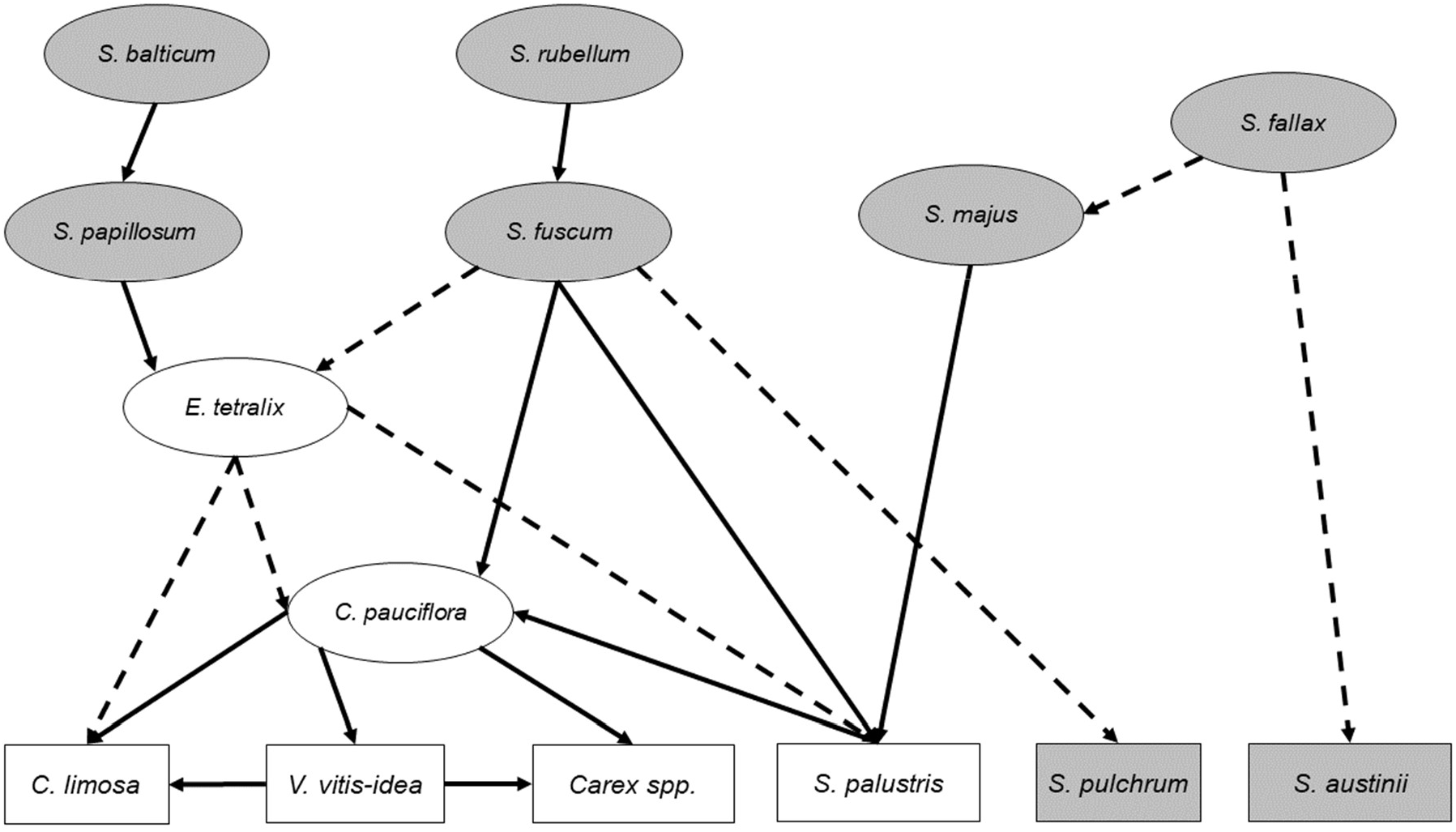
Graphical component of the partial Bayesian network that only includes strong interspecific relationships between predictor species (ovals) and co-responsive species (rectangles). Solid lines represent positive co-responses and dashed lines represent negative coresponses. *Sphagnum* mosses are shaded gray.

## DISCUSSION

We showed that modelling environmental co-responses among species from a European peat bog community improved the predictions of rare species. Based on a BN for the peat bog community, we identified two groups of species: co-responsive species that are typically rare and whose occurrence depends on the presence or absence of other species in the community, and predictor species that are more common and can be used to improve predictions of rare species. We analyzed a partial BN of only co-responsive and predictor species and found that this highly connected sub-network accounts for almost all the performance of the original BN. This finding suggests that only a small fraction of species and interspecific relationships, particularly those involving predictor species, need to be sampled to improve predictions of multiple rare species in an ecological community using our approach.

### Comparison of models

Notably, AUC scores for the eGLM+BN with 25% training data were similar to AUC scores for the eGLM with 50% training data (this trend is also apparent when comparing the eGLM+BN with 50% training data to the eGLM with 75% training data). This result suggests that using a BN to predict species occurrences can dramatically reduce the amount of data collection required to calibrate models. If information on environmental co-responses among species is available or can be estimated, then the eGLM+BN represents a viable method for improving the accuracy of species occurrence and community composition predictions, while adding minimal effort to the standard approach of environment-only models. The sGLM can also be used to reduce data collection, but it lacks some advantages of the eGLM+BN, described below.

The sGLM performed slightly better than the eGLM+BN when more data were available to calibrate models (i.e., with 75% training data). This result may be because the amount of information contained in a BN does not change as sample size changes, while parameters in the sGLM, particularly those involving non-focal species, are able to take advantage of the extra data for calibration. Or put another way: the BN summarizes a large amount of information about environmental co-responses in a standalone and compact form, which is why it can generate accurate occurrence predictions even with small amounts of calibration data. By contrast, at small sample sizes, the sGLM can misinterpret noise in data as a legitimate trend, resulting in less accurate predictions.

The difference between the sGLM and eGLM+BN is most prominent with rare species, whose environment-only model parameters may be especially unreliable due to the difficulty in finding locations at which they are known to be present. The sGLM is likely more sensitive to this unreliability because the effects of other species on the focal species are modelled as additional variables in a GLM, meaning that the benefits afforded by this extra information may remain overwhelmed by the baseline poor performance resulting from the bioclimate variables. By contrast, the eGLM+BN separates the modelling into an environment component (the eGLM part) and an interspecific component (the BN part)—for rare species and limited data, the information in the BN component can dominate the unreliable environment component, leading to comparatively better predictions.

The exceptional performance of the JSDM-inspired approach was unsurprising given the amount of information that can be incorporated in this model. However, to achieve this level of performance, a lot of empirical data is required to parameterize a complete and fully quantified correlation matrix. By contrast, the sGLM only requires knowledge of which species affect the presence of a focal species. Similarly, the eGLM+BN only requires knowledge on the presence of important interspecific relationships and the sign—positive or negative—of their effects (see Table S3 of Appendix S1 for a summary of data requirements for each model). Although using a Bayesian network with our simple assumptions about conditional dependencies can sometimes lead to unrealistic conditional probabilities (i.e., a probability of occurrence of 1 or 0 given the presence or absence of another species), such assumptions are unavoidable in a model that seeks to use as little data as the eGLM+BN. Besides the potential for the model to incorporate greater biological realism (which would hopefully reduce the frequency of these extreme predictions), discussed below, we argue that some lack of realism is permissible from a practical standpoint because it results in improved predictions compared to the eGLM (and for rare species and when data are limited compared to the sGLM). In many ways, it is remarkable that the eGLM+BN and sGLM get as close as they do to the performance of the JSDM-inspired approach. Overall, we consider the models in this study as offering a range of options to inform conservation decision-making.

Although the improved predictions produced by the eGLM+BN and sGLM both result from modelling interspecific relationships, each model may be better suited to describing different types of interspecific relationship. This expectation is realized as differences between the two models in species-level improvements in AUC over the eGLM (Appendix S1: Table S6). Some pairs of species may simply occur in a similar set of locations due to shared habitat preferences (or in a mutually exclusive manner due to different habitat preferences) in ways that are not described by the particular bioclimate variables included in the eGLM. In other words, we could attribute some predictive improvement resulting from multi-species models to more selective, hard-to-identify habitat preferences that are shared between species. The sGLM, which models the presences of other species in a similar way to bioclimate variables, should perform better when the set of non-focal species used in the model are known to have shared habitat preferences. Conversely, some co-occurrence relationships may be a result of biotic interactions, such as mutualism, competition or commensalism. Because effects of biotic interactions are less tied to bioclimate variables than shared habitat preferences, the eGLM+BN should perform better in these cases.

### Interpreting environmental co-responses among species

*Sphagnum* mosses are essential to the makeup of peat bog habitats because of the role species in this genus have as ecosystem engineers^48^. These mosses alter the composition of the soil in which they grow to reduce competition with other plants and increase their intake of nutrients and sunlight. This ability to modify the soil content of peat bogs makes *Sphagnum* mosses prime candidates for predictor species. Indeed, even though *Sphagnum* mosses made up only 37.0% of species from the peat bog community, six of the eight predictor species we identified were *Sphagnum* mosses, including *Sphagnum fuscum*, which is a dominant competitor of vascular plants^49^.

Although boosted regression tree analysis did not identify a relationship between *Sphagnum* classification and ΔAUC, *Sphagnum* mosses had an average ΔAUC of 0.048 compared to the *non-Sphagnum* average of 0.035, and 2 of the 6 co-responsive species we identified were *Sphagnum* mosses. These less common *Sphagnum* mosses often have very selective microhabitat preferences^50^, and to satisfy these preferences, they modify their habitats. But because many other plants cannot grow in the anoxic, low-nutrient soil favored by *Sphagnum* mosses, the presence of certain vascular plant species can be used as a signal for the absence of rare *Sphagnum* mosses.

### Adapting our approach to other ecological communities and for conservation

For other ecological communities, improving occurrence predictions using our approach would start by selecting a target species or set of species of interest. The next step is to determine which interspecific relationships involving the target species are worth modelling. We suggest two possible options: environmental co-responses and biotic interactions. As we did here, positive and negative relationships among species could be measured or estimated to identify a set of candidate species whose occurrences are strongly correlated with the target species. Alternatively, a set of candidate species could be developed based on which species have biotic interactions (e.g., competitive, facilitative) with the target species^17^. The set of candidate species from either option could then be refined by prioritizing species that fit the criteria for predictor species (i.e., species that are relatively common and insensitive to the presence of other species in the community) for inclusion in a BN. Environment-only models for these predictor and target species could then be combined with the streamlined BN to generate accurate predictions of the target species.

Predictor species appear to be a distinct group from umbrella and indicator species (Figure 3), making them a useful new category of conservation surrogate. As with all conservation surrogates, some initial analysis is necessary to identify these groups in a new ecological community^32^, but once identified, each group offers distinct benefits for particular aims. Umbrella species are characterized by their occurrence in a wide range of habitats^38^, and are used as conservation surrogates because their distributions often overlap with other species of interest^51^. However, umbrella species are often so widespread that relying on them to identify occurrences of rare species would lead to many false positives^40^. Indicator species are characterized by their occurrence in areas with high species richness^39^, and are used as conservation surrogates because their presence highlights locations with suitable conditions for a wide variety of species^52^. However, their presence is not guaranteed to inform the presence of rare species, which may have very different habitat preferences from more common species in the community^12^. Umbrella and indicator species offer a broad overview of ecosystem health and functioning to conservation practitioners^53,54^. Predictor species, which are defined by their relationship to rare species, offer a more detailed and finely resolved perspective that can complement umbrella and indicator species as part of a comprehensive conservation strategy.

**Figure 3.**
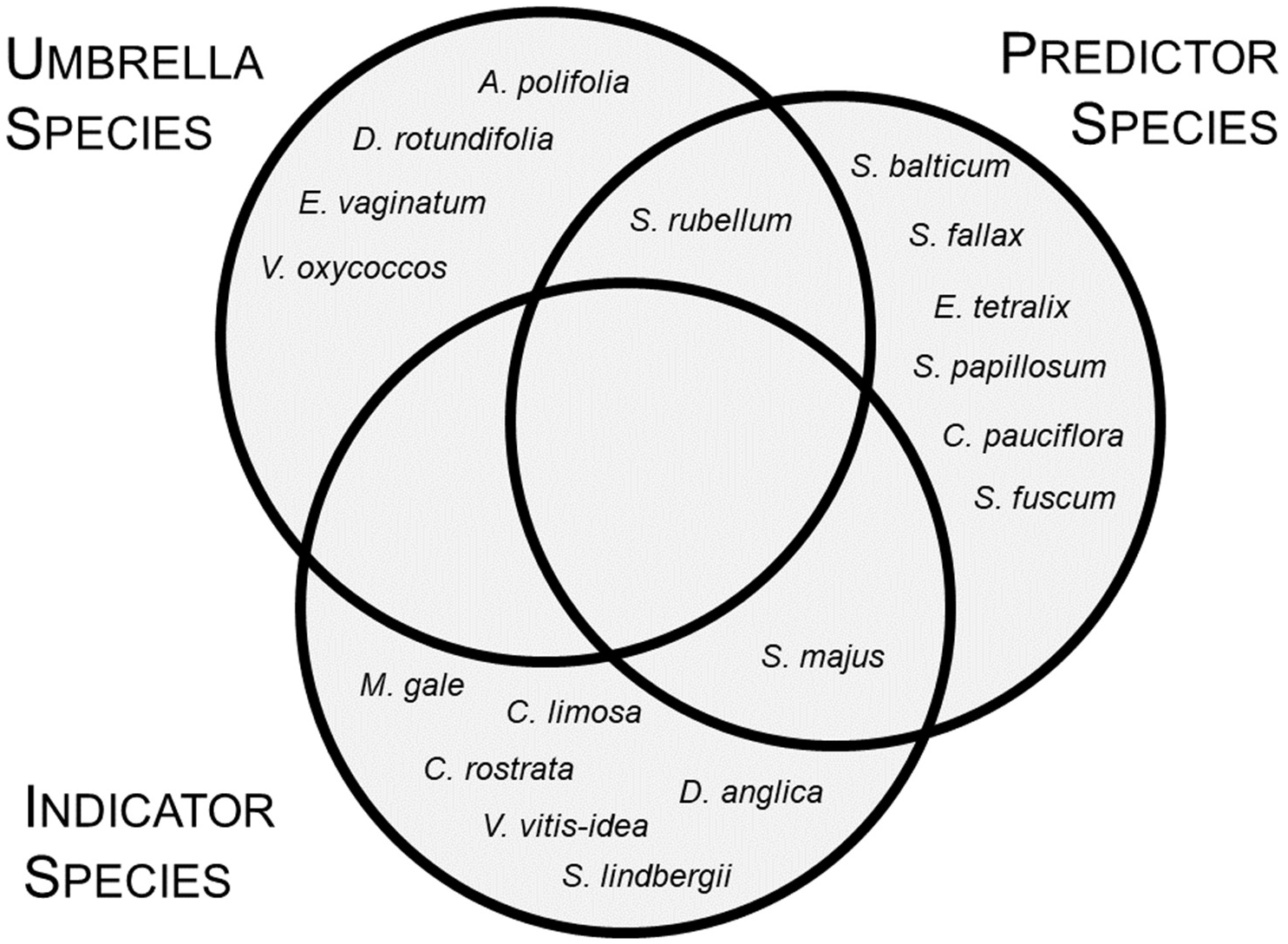
Venn diagram summarizing the overlap between umbrella, indicator, and predictor species. Notice that each group has a similar number of species but very few species belong to more than one group.

### Code availability

R code to run models and assess their performance using AUC will be made available on the Dryad digital repository if this manuscript is accepted. All data on the peat bog community are publicly available as part of a previously published study^34^.

## Acknowledgements

We thank Heather Lynch for comments on an early draft of this manuscript. P.R.T. was supported by a Maryland Summer Scholar Award and P.P.A.S. by a Postdoctoral Fellowship from the National Socio-Environmental Synthesis Center (SESYNC) funded by the National Science Foundation DBI-I052875.

## Author contributions

W.F. and P.P.A.S. designed the study; P.R.T. wrote code and performed analysis; P.R.T. wrote the first draft and all authors edited the manuscript; all authors have approved the final version of the manuscript.

