## Supplementary Figures, Tables and Methods for "Predictor species: Improving assessments of rare species occurrence by modelling environmental co-responses"

[3]Present address: Department of Biology, Brooklyn College, City University of New York, New York, NY, USA

This document consists of four figures, six tables, and an additional description of methods.

### SUPPLEMENTARY METHODS

#### Modifying data for Bayesian network analysis

We modified raw plant abundance data from peat bogs in Europe to use for generalized linear modelling, which requires the response variable to be on a binomial scale[40]. We acquired raw data in the form of two files, the first being a file containing abundance values for 54 plant taxa at 56 peat bog locations[36]. The authors of the study from which we obtain our data grouped together rare and closely related species (e.g., rare members of the *Sphagnum cuspidatum* subgenus) into one taxon to lower the number of very sparse (i.e., only present at one location) taxa[36]. We based all our occurrence prediction models on generalized linear models (GLMs), which require the choice of a specific link function; since we chose a logit link, we needed all the species occurrence data to be on a binomial ([0,1]) scale.

We were also given a data table containing bioclimate data for each of the European peat bog locations[36], which we used to calibrate our GLMs. Eleven variables were available for use in models: altitude, longitude, latitude, mean annual temperature, mean annual precipitation, temperature seasonality, precipitation seasonality, atmospheric deposition of sulfur, oxidized and reduced nitrogen, and Lang's moisture index, which is the ratio of precipitation to temperature in the warmest month in the year[36]. We limited our set of independent variables to those that are strongly related to the habitat preferences of plants. We identified which variables should be used in these models by computing Pearson correlation between each variable and species presence or absence (Table S1). We used mean annual temperature, latitude, mean annual precipitation, and temperature seasonality because these variables had relatively strong correlations with the presence or absence of the plant species in the peat bog community.

The bioclimate variables we chose were not highly correlated with each other, indicating that all of them provide unique value to the regression; none of the values had a correlation with absolute value above 0.55 (Table S2). We suggest that this result is likely because we used data from a study in Europe, an area with a variety of different climates (the Mediterranean region of Europe is quite warm relative to other areas with similar latitudes) and some surrounding islands (the British Isles, like most islands, likely have different climatic conditions than the nearby mainland).

#### Generalized linear models

We used R to generate GLMs for each plant species based on a set of random training partitions of locations[41]. The R "glm" function allows us to create individual GLMs for each species based on a subset of the 56 peat bog locations, and use these GLMs to predict the occurrence of these species at the rest of the locations. We used a data table containing the presence or absence of a species at each location, along with the four bioclimate variables, as data for GLMs. We used the "predict" function on a set of numeric entries representing the bioclimatic conditions at each location in the test partition. The result was a vector of probabilities for each species at all the locations in the test partition, allowing us to compare probabilities from our GLMs to other models as well as to themselves.

#### Using a BN to identify environmental co-responses among plant species

We created a binary, symmetric correlation matrix to use as a basis for the BN. We evaluated Pearson correlation values for each combination of plant species, and displayed the results in a similar, 54 x 54 matrix. We analyzed the effect that different correlation thresholds would have on the number of edges in the resulting BN to choose the optimal threshold (Figure S1). The finalized correlation matrix exclusively contains entries of 0, 1 and -1. With a large number of entries eliminated entirely, and any significant correlations rendered as equal, this resulting output is much more simple than a true correlation matrix.

Because the correlation matrix is symmetric and a BN cannot be, we invoked a hierarchy that directed the co-responses in the correlation matrix. Our hierarchy was based on the total abundance of each species over all 56 locations; with this hierarchy, individual species can only have incoming BN edges pointing from more abundant species. While we understand that this is likely not the nature of every interspecific interaction in the community, this hierarchy works generally well for a community that we know little about beyond our basic abundance data.

We evaluated eGLM+BN probabilities using the Boolean "OR" rule, which follows the following formula for a species $i$ (Figure S3):

$$p_{i,j}^{*} = \begin{cases} p_{i,j} + \min(p_{i,j}, 1 - p_{i,j}) & \text{if } n_{+,i,j} > n_{-,i,j} \\ p_{i,j} & \text{if } n_{+,i,j} = n_{-,i,j} \\ p_{i,j} - \min(p_{i,j}, 1 - p_{i,j}) & \text{if } n_{+,i,j} < n_{-,i,j} \end{cases}$$

where $p_{i,j}^{*}$ is the posterior probability of species $i$ at location $j$, and $p_{i,j}$ is the prior probability of species $i$ at location $j$; $n_{+,i,j}$ and $n_{-,i,j}$ are the number of species with positive and negative BN edges pointing to species $i$ that are present at location $j$, respectively[17].

Using this rule of thumb, we can develop a conditional probability table given prior probabilities for each species and a BN. Consider a community where Species A, B, and C have priors 0.8, 0.3, and 0.6 respectively (Figure S3). If we are given a BN that includes two edges, one positive edge pointing from A to C and one negative edge pointing from B to C, we can use a conditional probability table to more accurately estimate the true presence rate of C. Because C has two incoming BN edges, we generate a conditional probability table for the 4 ($2^2$) possible combinations of presence for Species A and B (A present only, B present only, both present or both absent). For each of these hypothetical scenarios we can calculate $n_{+,C}$ and $n_{-,C}$. When Species A is present and Species B is not, $n_{+,C}$ = 1 and $n_{-,C}$ = 0, so the corresponding entry of the conditional probability table is $p_C + \min(p_C, 1 - p_C)$, where $p_C$ is 0.6 as previously stated. This results in the first entry of the conditional probability table (corresponding to situations where A is present and B is absent) being 1. The other three entries are calculated similarly using the above equation for the posterior probability, and then once the conditional probability table is complete, the revised posteriors are calculated using Bayes' formula.

**Developing a joint species distribution model to quantify co-occurrence relationships**

We used joint multivariate logistic regression to produce estimates for a species' occurrence at a given location using a full correlation matrix for the plant community[34]. We interpreted the presences or absences of each plant in the peat bog community as one component of a normally distributed random vector (a vector where each component is normally distributed around a corresponding component of a mean vector, and each component may depend on the others based on a correlation matrix). Given a mean vector and a correlation matrix, one can obtain random values and analyze their correlation in order to obtain information about interspecific relationships[34,35]; however, we used this distribution differently because we assumed a known presence or absence for every non-focal species in our analysis. We obtained the conditional distribution of one component of the multivariate vector given the values of the other components (this conditional distribution is normal), using this distribution to estimate the probability that a species was present. We developed the mean vector for this distribution based on the eGLM predictions for each species[34], so the revised conditional distribution given the other values of the components was shifted according to the presence or absence of the other species.

**Interpreting AUC scores from the three occurrence prediction models**

We used the area under the receiver operating characteristic curve (AUC) method to evaluate the predictive accuracy of each model. AUC scores range from 0 to 1; a score of 1 represents a perfectly correct prediction, while a score of 0.5 represents random guessing[42]. It is then very much expected for all AUC scores in our analysis to be greater than 0.5, and given our findings with our JSDM-inspired approach, we cannot expect AUC scores to be greater than 0.85 for more simple models such as the eGLM, sGLM or eGLM+BN. Therefore, a difference in AUC between models of 0.1 or higher is very significant[42]. Therefore, we choose 0.08 as the ΔAUC value that a species needs to consistently attained for consideration as a co-responsive species (Table S4).

We also experimented with true skill statistic (TSS)[43], as an alternative to AUC for model evaluation. Unlike AUC, TSS values range from -1 to 1, with 0 representing random guessing. We found that while in some cases TSS and AUC produced similar results, the sGLM outperforms the eGLM+BN at all training sizes when we used TSS to evaluate model performance (Figure S4). TSS values were also much more variable from one randomization to another, making it difficult to confirm the significance of any trends we found. For some random partitions of the 56 locations, the average TSS for each species was close to 0.1 for all three models, while other times this average was close to 0.5. With variability this high it is difficult to draw conclusions or identify various trends as statistically significant.

**Identifying which traits were most pivotal in identifying co-responsive species**

Boosted regression tree (BRT) analysis is a prediction method consisting of the boosting of a large set of decision trees. Decision trees, which predict the value of a dependent variable by checking the value of many independent variables in relation to defined cut-off points[46], are a basic but effective prediction tactic. However, single decision trees are not very smooth, and the cut-off points can be chosen rather arbitrarily; BRT analysis uses many trees and takes a weighted average of their outputs[45]. Individual trees that are more accurate are given higher weights. When hundreds or thousands of trees are incorporated, the prediction becomes much smoother. We used 1,000 trees in our analysis and trained the model randomly with 75% of the data. Literature on BRT analysis recommends smaller training fractions[45] but because we only had 54 data points, we could not obtain BRT predictions using a smaller value. We calibrated, tested and evaluated BRT models using the R "gbm" package with little computational expense.

BRT analysis suggests that rarity is very important in determining whether a species is likely to have a high ΔAUC. We explored whether this trend may simply be a consequence of the abundance-based hierarchy of influence we used to construct the BN, which prevented highly abundant species from having incoming BN edges. Although individual species abundance across all locations was positively correlated with rarity ($r = 0.741$, $p < 0.001$), BRT analysis showed that rarity still had a much stronger relationship with ΔAUC than abundance or, more importantly, the number of incoming BN edges. If rare species had high ΔAUC values only because they were on the bottom of the hierarchy, any species with many incoming BN edges (or any incoming BN edges, for that matter) would have a high ΔAUC, but this is not the case.

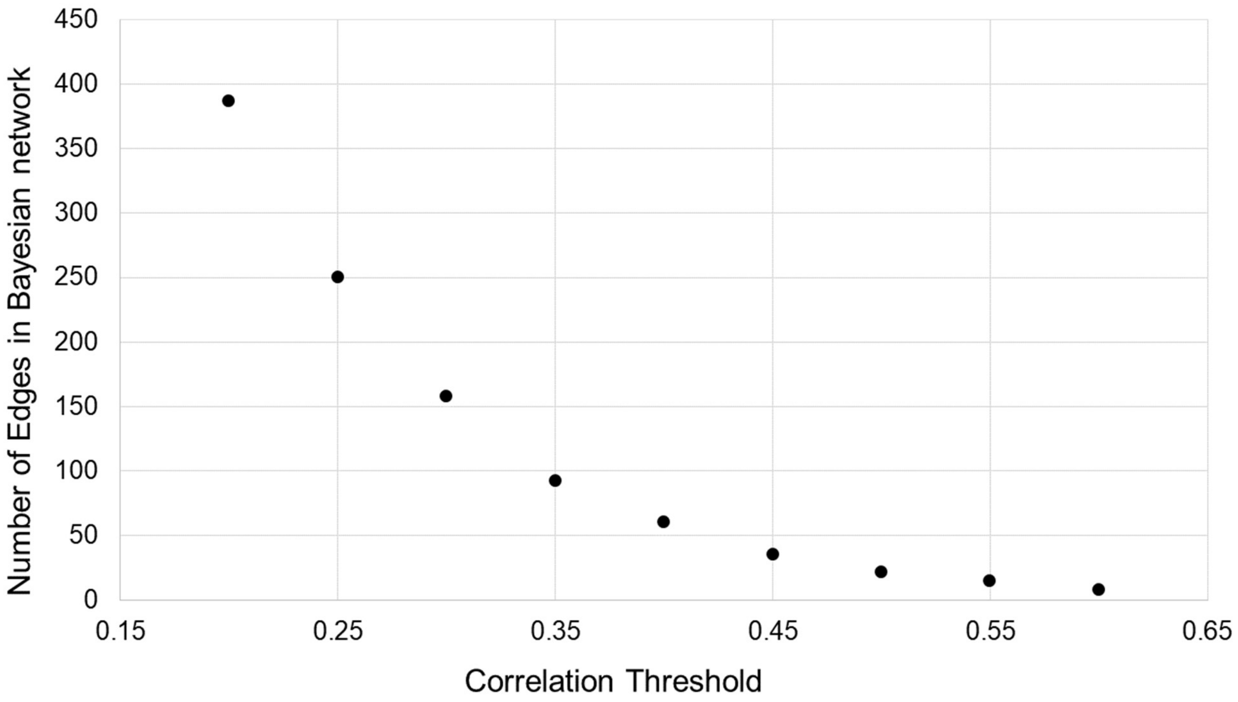


**SUPPLEMENTARY FIGURE 1**

**Supplementary Figure 1 – Number of edges in a resulting Bayesian network (BN) based on correlation threshold.** Note that a correlation threshold of 0.35 represents a clear point of inflection; values above 0.35 result in a BN that would leave many species without any edges, while values below 0.35 result in a BN that would potentially contain too many frivolous edges.

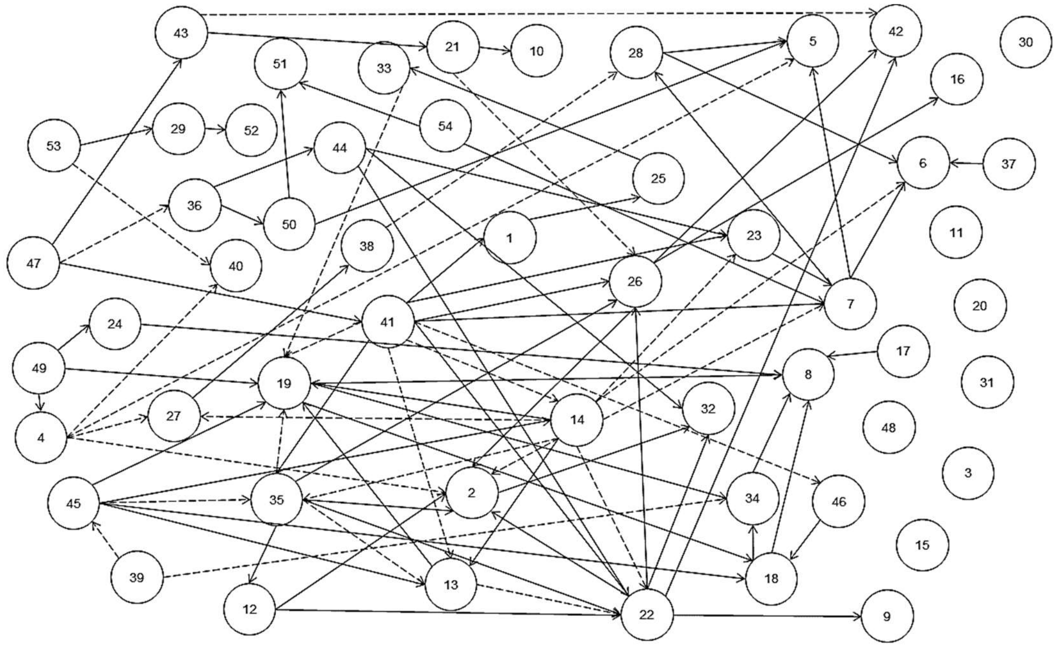


1 *Andromeda polifolia*
2 *Betula nana*
3 *Betula pubescens*
4 *Calluna vulgaris*
5 *Carex spp*
6 *Carex limosa*
7 *Carex pauciflora*
8 *Carex rostrata*
9 *Drosera anglica*
10 *Drosera intermedia*
11 *Drosera rotundifolia*
12 *Empetrum nigrum*
13 *Eriophorum angustifolium*
14 *Erica tetralix*
15 *Eriophorum vaginatum*
16 *Ledum palustre*
17 *Molinea caerulea*
18 *Myrica gale*
19 *Narthecium ossifragum*
20 *Pinus sylvestris*
21 *Rhynchospora alba*
22 *Rubus chamaemorus*
23 *Scheuchzeria palustris*
24 *Trichophorum cespitosum*
25 *Vaccimium oxycoccos*
26 *Vaccinium uliginosum*
27 *Vaccinium microcarpon*
28 *Vaccinium vitis-idea*
29 *Cladonia spp*
30 *Sphagnum section Acutifolia*
31 *Sphagnum section Sphagnum*
32 *Sphagnum section Cuspidata*
33 *Sphagnum angustifolium*
34 *Sphagnum austinii*
35 *Sphagnum balticum*
36 *Sphagnum capillifolium*
37 *Sphagnum compactum*
38 *Sphagnum cuspidatum*
39 *Sphagnum fallax*
40 *Sphagnum flexuosum*
41 *Sphagnum fuscum*
42 *Sphagnum lindbergii*
43 *Sphagnum magellanicum*
44 *Sphagnum majus*
45 *Sphagnum papillosum*
46 *Sphagnum pulchrum*
47 *Sphagnum rubellum*
48 *Sphagnum russowii*
49 *Sphagnum tenellum*
50 *Bryales*
51 *Funariales*
52 *Dicranales*
53 *Hypnales*
54 *Polytrichales*

### SUPPLEMENTARY FIGURE 2

**Supplementary Figure 2 – The full Bayesian network (BN) produced for our peat bog community.** All but seven species (located on the right side of the graph) are included. Continuous lines represent positive edges, while dashed lines represent negative edges. The entire BN has 92 edges in total; 65 are positive and 27 are negative. We also include a legend displaying the names of each species corresponding to numbers on the graph.

a.

| Species | eGLM Probability |
|---|---|
| A | 0.8 |
| B | 0.3 |
| C | 0.6 |
| D | 0.5 |

b.

| | A | B | C | D |
|---|---|---|---|---|
| A | 0 | 0 | 1 | 0 |
| B | 0 | 0 | -1 | 0 |
| C | 0 | 0 | 0 | 0 |
| D | 0 | 0 | 0 | 0 |

c.

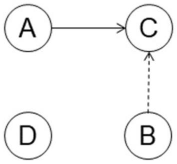


d.

| Sp. A | Sp. B | Formula | P(C) | |
|---|---|---|---|---|
| 1 | 0 | P(C) + min(P(C),1-P(C)) | 1.0 | $q_{C\|A=1,B=0} = 1.0$ |
| 0 | 0 | P(C) | 0.6 | $q_{C\|A=0,B=0} = 0.6$ |
| 1 | 1 | P(C) | 0.6 | $q_{C\|A=1,B=1} = 0.6$ |
| 0 | 1 | P(C) – min(P(C),1-P(C)) | 0.2 | $q_{C\|A=0,B=1} = 0.2$ |

$$q_C = q_{C|A=1,B=0} * p_A(1-p_B) + q_{C|A=0,B=0} * (1-p_A)(1-p_B) + q_{B|A=1,B=1} * p_A(p_B) + q_{C|A=0,B=1} * (1-p_A)(p_B)$$

$$q_C = 1.0 * 0.56 + 0.6 * 0.14 + 0.6 * 0.24 + 0.2 * 0.06 = 0.8$$

**eGLM+BN Probability for Species C: 0.8**

### SUPPLEMENTARY FIGURE 3

**Supplementary Figure 3 - Example workflow for calculating occurrence probabilities using a Bayesian network that represents environmental co-responses among a community of four species.** (a) “Prior” occurrence probabilities for four species at a particular location are first obtained from an eGLM, which takes into account only environmental conditions at a specific location. (b) Correlations between the occurrence of each pair of species at all sampled locations are used to identify strong positive (e.g., A and C) and negative (e.g., B and C) environmental co-responses among species. (c) A hierarchy of species (A above B, B above C, C above D) is used to determine the direction of each influence, described by the graphical component of the Bayesian network. (d) The second component of the Bayesian network is a conditional probability table for C that specifies how the occurrence of A and B at a location affects the occurrence probability of C, and, below, the calculation of the “posterior” occurrence probability for C at the example location, which now takes into account environmental co-responses as well as abiotic conditions. Notice that the probability for C with the eGLM+BN is higher than with the eGLM because the probability of A (positive co-response with C) being present at the location is higher than the probability of B (negative co-response with C).

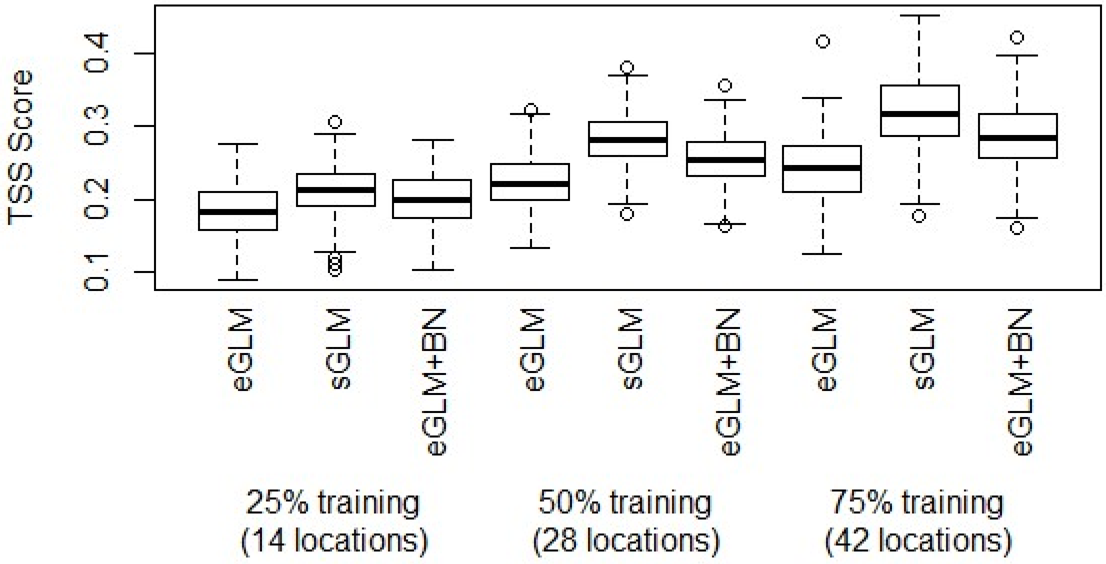


**SUPPLEMENTARY FIGURE 4**

**Supplementary Figure 4 - Performance of the eGLM, sGLM, and eGLM+BN measured by TSS at three training partition sizes.** While some trends regarding the three models are similar regardless of what evaluation tool is used, the sGLM performs better than the eGLM+BN at all training data sizes. TSS scores also vary much more than AUC scores from one randomization to another.

**SUPPLEMENTARY TABLE 1**

| Bioclimate Variable | Average Correlation with Species Presence/Absence |
|---|---|
| Temperature Seasonality | 0.260 |
| Latitude | 0.253 |
| Mean Annual Temperature | 0.243 |
| Mean Annual Precipitation | 0.200 |
| Longitude | 0.191 |
| Lang's moisture index | 0.173 |
| Precipitation Seasonality | 0.169 |
| Altitude | 0.169 |
| Reduced nitrogen concentration | 0.164 |
| Atmospheric sulfur concentration | 0.160 |
| Oxidized nitrogen concentration | 0.152 |

**Supplementary Table 1 – Absolute values of Pearson correlation between each bioclimate variable and the occurrence of species across all locations, on average (n = 56).** Underlined variables were used in generalized linear models (GLMs) because of their relatively high correlations.

### SUPPLEMENTARY TABLE 2

| **Bioclimate Variable** | Mean Annual Temperature | Mean Annual Precipitation | Temperature Seasonality | Latitude |
|---|---|---|---|---|
| Mean Annual Temperature | | 0.468 | -0.548 | -0.414 |
| Mean Annual Precipitation | | | 0.015 | 0.136 |
| Temperature Seasonality | | | | 0.475 |
| Latitude | | | | |

**Supplementary Table 2 – Table depicting correlations between all the bioclimate variables used in the eGLM and subsequent models**. None of these variables are correlated at an abnormally strong value (the highest being roughly 0.55), suggesting that there are no overfitting issues in the eGLM, sGLM or eGLM+BN. Obviously the variables have some sort of relationship, but since none of the correlations are remotely linear for any pair of bioclimate variables, each variable is important in the construction of GLMs for species occurrence predictions.

**SUPPLEMENTARY TABLE 3**

| Model | Data requirements |
|---|---|
| eGLM | • Environmental data on each location in the community<br>• Presences/absences for all species at locations in the training partition |
| sGLM | • Environmental data on each location in the community<br>• Presences/absences for all species at locations in the training partition<br>• Knowledge of any interspecific interactions with the focal species (co-occurrence relationships, biotic interactions, etc.)<br>• Presences/absences for species interacting with the focal species at all locations |
| eGLM+BN | • Environmental data on each location in the community<br>• Presences/absences for all species at locations in the training partition<br>• Knowledge of any interspecific interactions with the focal species (co-occurrence relationships, biotic interactions, etc.) and some knowledge about the nature of these interactions (positive/negative)<br>• Presences/absences for species interacting with the focal species at all locations |
| JSDM-inspired approach | • Environmental data on each location in the community<br>• Presences/absences for all species at locations in the training partition<br>• Presences/absences for all species at locations in the test partition<br>• Exact correlation values for each pair of species in the community (more than just a knowledge of interactions) |

**Supplementary Table 3 – Table describing the data requirements for each model.** Obviously the eGLM, which does not incorporate species interactions, requires the smallest amount of data, but for multi-species models, both the sGLM and eGLM+BN do not require very much additional data. The approach inspired by joint species distribution modelling (JSDM) techniques requires data that may be difficult or impossible to collect in some situations, making it a potentially unrealistic model choice. As a result, we focus on how the eGLM+BN or sGLM can be used to facilitate improved conservation practices with a much lower amount of input data.

### SUPPLEMENTARY TABLE 4

| Number of random sets, η | Number of species with ΔAUC > 0.08 in at least η sets |
|---|---|
| 1 | 10 |
| 2 | 9 |
| 3 | 8 |
| 4 | 8 |
| 5 | 8 |
| 6 | 7 |
| 7 | 6 |
| 8 | 6 |
| 9 | 6 |
| 10 | 3 |

**Supplementary Table 4 – Table describing the prevalence of high ΔAUC (change in AUC score average from eGLM to eGLM±BN at 50% training partition) values in the different random sets, while also explaining the variability in each set.** Species with ΔAUC above 0.08 in at least 9 of the 10 random sets (shaded in light gray) were classified as co-responsive species; we identified 6 co-responsive species in the peat bog plant community.

**SUPPLEMENTARY TABLE 5**

| Species | Rarity (%) | ΔAUC with partial BN | ΔAUC with original BN |
|---|---|---|---|
| *Carex spp.* | 8.93 | 0.124 ± 0.232 | 0.134 ± 0.178 |
| *Carex limosa* | 10.71 | 0.147 ± 0.141 | 0.099 ± 0.165 |
| *Vaccinium vitis-idea* | 9.72 | 0.130 ± 0.166 | 0.097 ± 0.153 |
| *Scheuchzeria palustris* | 17.86 | 0.128 ± 0.106 | 0.126 ± 0.137 |
| *Sphagnum austinii* | 8.93 | 0.199 ± 0.093 | 0.088 ± 0.174 |
| *Sphagnum pulchrum* | 12.50 | 0.153 ± 0.135 | 0.139 ± 0.114 |

**Supplementary Table 5 – ΔAUC values (mean ± standard deviation) for the six co-responsive species with partial and original Bayesian networks (BNs).** Note the similarity, and in many cases, improvement, between ΔAUC values for each species. We also include rarity values to display how co-responsive species generally occur at an exceptionally low proportion of the 56 peat bog sites.

### SUPPLEMENTARY TABLE 6

| Species Name | eGLM AUC | sGLM-eGLM AUC | ΔAUC |
|---|---|---|---|
| *Andromeda polifolia* | 0.785 ± 0.130 | 0.005 ± 0.114 | 0.026 ± 0.111 |
| *Betula nana* | 0.820 ± 0.099 | -0.110 ± 0.117 | 0.039 ± 0.127 |
| *Carex spp* | 0.632 ± 0.203 | 0.178 ± 0.223 | 0.134 ± 0.178 |
| *Carex limosa* | 0.690 ± 0.157 | 0.036 ± 0.196 | 0.099 ± 0.165 |
| *Carex pauciflora* | 0.803 ± 0.122 | 0.002 ± 0.141 | 0.029 ± 0.149 |
| *Carex rostrata* | 0.816 ± 0.216 | -0.017 ± 0.180 | -0.003 ± 0.169 |
| *Drosera anglica* | 0.653 ± 0.095 | 0.051 ± 0.113 | 0.044 ± 0.113 |
| *Drosera intermedia* | 0.642 ± 0.163 | 0.099 ± 0.134 | 0.067 ± 0.124 |
| *Empetrum nigrum* | 0.876 ± 0.077 | -0.004 ± 0.045 | -0.018 ± 0.095 |
| *Eriophorum angustifolium* | 0.826 ± 0.083 | -0.025 ± 0.102 | 0.046 ± 0.120 |
| *Erica tetralix* | 0.868 ± 0.079 | -0.036 ± 0.059 | 0.010 ± 0.109 |
| *Ledum palustre* | 0.861 ± 0.198 | -0.027 ± 0.154 | 0.030 ± 0.140 |
| *Myrica gale* | 0.871 ± 0.157 | 0.038 ± 0.126 | -0.004 ± 0.147 |
| *Narthecium ossifragum* | 0.909 ± 0.085 | -0.019 ± 0.114 | 0.000 ± 0.169 |
| *Rhynchospora alba* | 0.678 ± 0.080 | 0.039 ± 0.054 | 0.063 ± 0.082 |
| *Rubus chamaemorus* | 0.907 ± 0.080 | -0.076 ± 0.102 | 0.001 ± 0.130 |
| *Scheuchzeria palustris* | 0.618 ± 0.105 | 0.176 ± 0.145 | 0.126 ± 0.137 |
| *Trichophorum cespitosum* | 0.690 ± 0.095 | 0.066 ± 0.070 | 0.051 ± 0.099 |
| *Vaccimium oxycoccos* | 0.695 ± 0.110 | 0.021 ± 0.099 | 0.014 ± 0.103 |
| *Vaccinium uliginosum* | 0.803 ± 0.116 | -0.022 ± 0.124 | 0.035 ± 0.143 |
| *Vaccinium microcarpon* | 0.750 ± 0.142 | 0.005 ± 0.113 | -0.037 ± 0.125 |
| *Vaccinium vitis-idea* | 0.674 ± 0.169 | 0.076 ± 0.177 | 0.097 ± 0.153 |
| *Cladonia spp* | 0.675 ± 0.086 | 0.026 ± 0.058 | 0.036 ± 0.073 |
| *Sphagnum section Cuspidata* | 0.638 ± 0.238 | -0.012 ± 0.184 | 0.027 ± 0.163 |
| *Sphagnum angustifolium* | 0.674 ± 0.087 | 0.032 ± 0.070 | 0.038 ± 0.081 |
| *Sphagnum austinii* | 0.797 ± 0.196 | -0.022 ± 0.166 | 0.088 ± 0.174 |
| *Sphagnum balticum* | 0.966 ± 0.046 | -0.014 ± 0.060 | -0.001 ± 0.103 |
| *Sphagnum capillifolium* | 0.582 ± 0.079 | 0.064 ± 0.102 | 0.079 ± 0.096 |
| *Sphagnum compactum* | 0.824 ± 0.278 | -0.051 ± 0.152 | 0.014 ± 0.158 |
| *Sphagnum flexuosum* | 0.620 ± 0.146 | 0.062 ± 0.172 | 0.069 ± 0.137 |
| *Sphagnum fuscum* | 0.808 ± 0.146 | 0.004 ± 0.053 | -0.016 ± 0.077 |
| *Sphagnum lindbergii* | 0.803 ± 0.205 | 0.002 ± 0.171 | 0.038 ± 0.171 |
| *Sphagnum magellanicum* | 0.693 ± 0.099 | 0.058 ± 0.080 | 0.066 ± 0.104 |
| *Sphagnum majus* | 0.642 ± 0.093 | 0.029 ± 0.072 | 0.038 ± 0.091 |
| *Sphagnum papillosum* | 0.719 ± 0.084 | 0.000 ± 0.055 | 0.013 ± 0.076 |
| *Sphagnum pulchrum* | 0.583 ± 0.107 | 0.094 ± 0.124 | 0.139 ± 0.114 |
| *Sphagnum tenellum* | 0.583 ± 0.087 | 0.066 ± 0.100 | 0.086 ± 0.097 |
| *Bryales* | 0.592 ± 0.085 | 0.070 ± 0.148 | 0.005 ± 0.112 |
| *Funariales* | 0.627 ± 0.124 | 0.107 ± 0.170 | 0.008 ± 0.152 |
| *Dicranales* | 0.640 ± 0.080 | 0.035 ± 0.070 | 0.019 ± 0.078 |

**Supplementary Table 6 – eGLM AUC values, ΔAUC values and (sGLM-eGLM) AUC values (mean ± standard deviation) for every taxon in the peat bog community that had incoming BN edges (i.e., cases where the multi-species models could be used to potentially improve eGLM predictions) at the 50% training partition.** This data comes from 1,000 random partitions of the 56 locations in the peat bog community. Co-responsive species are shaded gray.
